# SpiceRx: an integrated resource for the health impacts of culinary spices and herbs

**DOI:** 10.1101/273599

**Authors:** Rakhi Nk, Rudraksh Tuwani, Neelansh Garg, Jagriti Mukherjee, Ganesh Bagler

## Abstract

Spices and herbs are key dietary ingredients used in cuisines across the world. They have been reported to be of medicinal value for a wide variety of diseases through a large body of biomedical investigations. Bioactive phytochemicals in these plant products form the basis of their therapeutic potential as well as adverse effects. A systematic compilation of empirical data involving these aspects of culinary spices and herbs could help unravel molecular mechanisms underlying their effects on health.

SpiceRx provides a platform for exploring the health impact of spices and herbs used in food preparations through a structured database of tripartite relationships with their phytochemicals and disease associations. Starting with an extensive dictionary of culinary spices and herbs, their disease associations were text mined from MEDLINE, the largest database of biomedical abstracts, assisted with manual curation. This information was further combined with spice-phytochemical and phytochemical-disease associations. SpiceRx is an integrated repertoire of evidence-based knowledge pertaining to the health impacts of culinary spices and herbs, and facilitates their disease-specific culinary recommendations as well as exploration of molecular mechanisms underlying their health effects.

**Availability and Implementation:** SpiceRx is available at http://cosylab.iiitd.edu.in/spicerx and supports all modern browsers. SpiceRx is implemented with Python web development framework Django and relational database PostgreSQL; the front-end was built using HTML, CSS, JavaScript, AJAX, jQuery, JSME Molecular Editor, Bootstrap, Jmol, DataTables and Google Charts.

**Supplementary information:** Supplementary data are available at *Bioinformatics* online.

## Introduction

Spices and herbs have a unique place in culinary preparations across the world cuisines (Singh and Bagler, unpublished; CulinaryDB, http://cosylab.iiitd.edu.in/culinarydb). While they have been appreciated as flavoring agents and suggested to be of value as antimicrobial agents, a coherent picture of their health effects is hitherto unavailable (Billing and Sherman, 1998). Reductionist investigations have primarily focused on benevolent and adverse effects of individual spices or their phytochemicals, presenting a potpourri of facts with no coherent picture (Srinivasan, 2005b; Yashin *et al.*, 2017; Srinivasan, 2005a). While such studies have added to the fragments of evidence at the levels of culinary spices/herbs and their molecular constituents, a holistic understanding of their health impacts remains vague and unstructured.

Integration of scientific evidence available from an exponentially growing literature reporting health consequences of culinary spices and herbs will enable drawing inferences for their informed culinary use as well as for generating hypotheses to discover underlying molecular mechanisms (Rakhi *et al.*, unpublished). SpiceRx bridges information associating culinary spices/herbs, their phytochemicals and diseases with the help of evidence compiled from research articles and external resources to provide a platform for open-ended explorations of their tripartite relationships (Figure 1).

**Figure 1:**
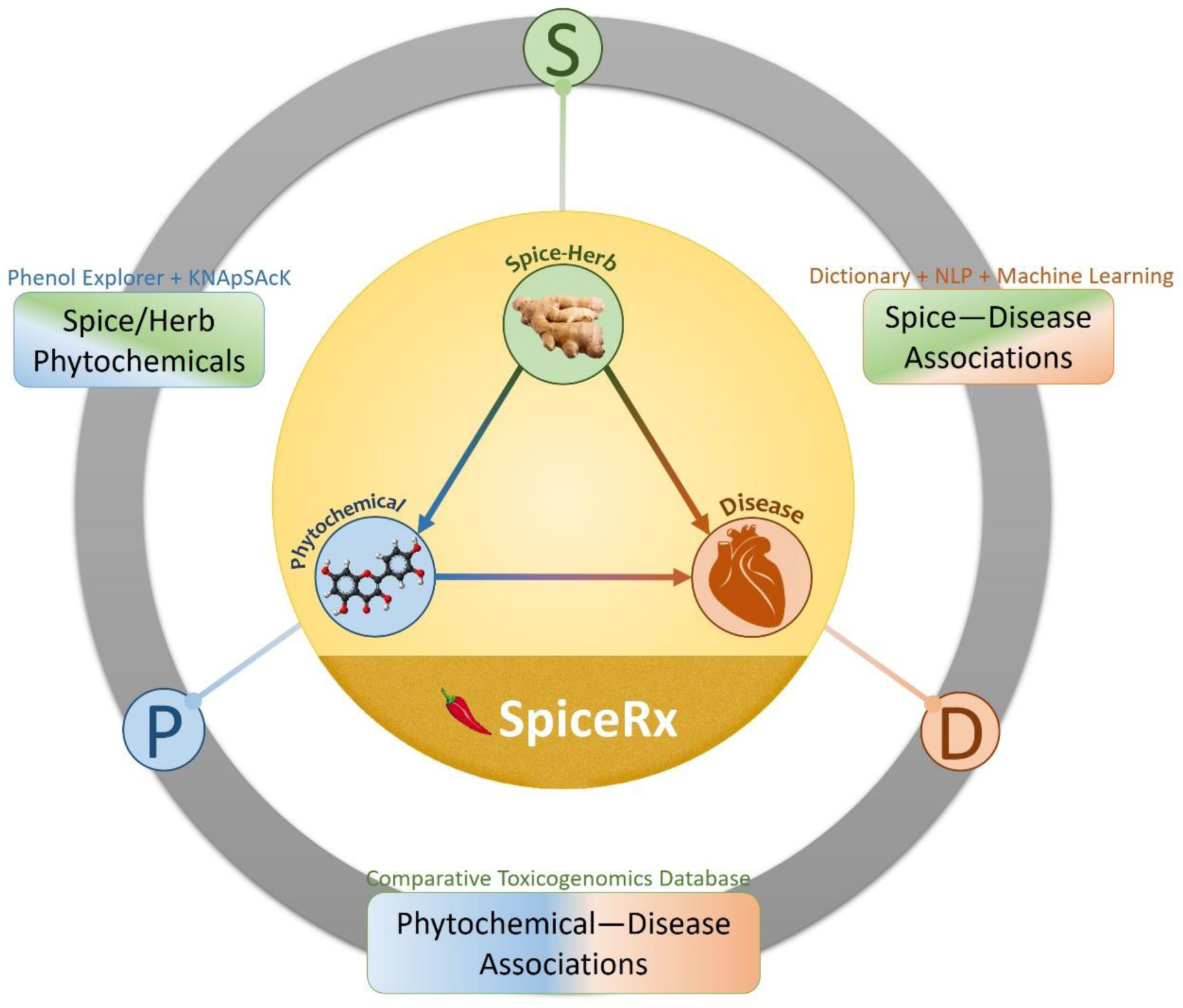
SpiceRx is an integrated repertoire of evidence-based knowledge pertaining to the health impacts of culinary spices/herbs and their phytochemicals. It seamlessly integrates scientific evidence from biomedical literature and external resources using a text mining protocol.

## Database Overview

SpiceRx provides a platform for exploring the health impact of spices and herbs used in food preparations through a structured database of tripartite relationships with their phytochemicals and disease associations (Figure 1). Starting with an extensive dictionary of 188 culinary spices and herbs, their disease associations were text mined from 28 million MEDLINE abstracts. These data were further combined with evidence of spice-phytochemical and phytochemical-disease associations. SpiceRx presents a compilation of 11750 MEDLINE abstracts containing 8957 disease associations (8172 positive and 783 negative) for 152 spices linked to 848 unique disease-specific Medical Subject Headings (MeSH) IDs (Lipscomb, 2000). The hierarchical organization of MeSH was used as a basis for ontological classification of disease terms. Spice names were tagged using a dictionary matching method whereas diseases were tagged using a machine learning based model, TaggerOne (Leaman and Lu, 2016). A convolutional neural network based relation extraction model (Nguyen and Grishman, 2015; Kumar Sahu *et al.*, 2016) was trained on 6712 manually annotated sentences to extract and classify positive, negative and neutral associations.

Information of 866 phytochemicals was obtained for 142 of the spices using KNApSAcK (Afendi *et al.*, 2012) and PhenolExplorer (Rothwell *et al.*, 2013) that comprised of 570 bioactive compounds with a total of 2042 spice-phytochemical associations. These compounds were further linked to diseases with the help of Comparative Toxicogenomics Database (CTD) (Davis *et al.*, 2017), a public database of literature-curated and inferred chemical-disease associations. The resource unearths literature-supported spice-disease associations for which the spice phytochemical(s) have been independently reported for therapeutic effects. Thus, through interlinked triangular relationships between culinary herbs and spices, their phytochemicals, and diseases, SpiceRx facilitates seamless exploration of evidence-based knowledge for their disease-specific culinary recommendations as well as enquiry into underlying molecular mechanisms.

## Architecture and Web Interface

SpiceRx has been designed to facilitate explorations starting from either a spice/herb, a disease, or a phytochemical query to find its association with the remaining two elements.

### Spice Search

Culinary spices/herbs can be searched using their common name, scientific name or NCBI taxonomy ID to obtain their disease associations and constituent phytochemicals. Searching for a spice yields paginated list of its disease associations (ranked in descending order of number of publications) and that of its phytochemicals. Research articles reporting the association are listed with link-outs to PubMed. For 20% associations, therapeutic spice phytochemicals involved in a spice-disease association discovered through triangular causal linking are listed. These provide a ground for their pharmaceutical and nutraceutical applications in addition to informed culinary use. Separately, a list of all spice-linked phytochemicals is provided. Phytochemicals could be further explored for their detailed physicochemical features, drug-likeness, and links to associated spices and diseases.

### Disease Search

To identify culinary spices and herbs that are reported with either therapeutic or adverse effect against a specific disease, SpiceRx provides disease search integrated with the hierarchical organization of MeSH disease terms. One may search by Disease Name, Disease Category, Disease Sub Category or MeSH ID. The Disease Name could be used to search by common names, such as ‘Diabetes‘, ‘Weight Gain‘, or ‘Obesity‘. The ‘MeSH Disease Category’ represents a broad class of diseases such as ‘Cardiovascular Diseases‘, ‘Endocrine System Diseases’ or ‘Neoplasms‘, whereas ‘Disease Sub Category’ represents more refined disease terms such as ‘Heart Diseases‘, ‘Metabolic Diseases‘, ‘Liver Diseases’ etc. A null search performed with no specific disease term is designed to present an exhaustive list of diseases and their spice associations. Details of spices reported with positive or negative effects for the queried disease term along with link-outs to PubMed articles are presented, in addition to any specific phytochemical(s) involved in the therapeutic association, whenever available.

### Phytochemical Search

Spice phytochemicals form the basis for molecular mechanisms involved in therapeutic effects against specific diseases. Apart from querying SpiceRx for a spice or a disease, one may also explore the resource to search compounds by their structure, common name, IUPAC name, PubChem ID, molecular weight, Hydrogen bond donors/acceptors or molecular hydrophobicity (AlogP). Each phytochemical could further be explored for its chemical profile (MESH ID, PubChem ID, Common name, IUPAC name, Molecular Formula, Canonical and Isomeric SMILES; Phytochemical and ADMET properties), disease associations as well as a list of spices in which it is reported to be found in. Apart from 2D and 3D visualizations and download options (Mol, 2D Image, and SDF), lookup for structurally similar spice compounds within the database as well as those commercially available from external sources is also provided. The ‘null search’ (with all fields empty) is designed to yield a list of all spice phytochemicals.

Please see Supplementary Information for details of materials and methods, tech stack, database statistics, web interface search features, use cases and data download options.

## Conclusions and Discussion

SpiceRx integrates scientific evidence from biomedical literature and external resources to seamlessly collate tripartite associations between culinary spices/herbs, diseases, and phytochemicals. By blending scattered and disorganized evidence, it provides a platform for investigation of spices for their health effects, chemical mechanisms behind their action, and paves way for developing nutraceuticals and drugs.

The effect of spice/herb on a disease may sensitively depends on its quantity. Due to unavailability of uniform data, SpiceRx does not include this information. Further, disease associations can vary according various factors such as the age, pre-existing conditions and gender among others, which are not represented at present. There is scope to enhance SpiceRx to include such additional features.

## Acknowledgements

G.B. thanks the Indraprastha Institute of Information Technology (IIIT-Delhi) for providing computational facilities and support. R.N.K. thanks the Ministry of Human Resource Development, Government of India and Indian Institute of Technology Jodhpur for the senior research fellowship. R.T. (Research Associate), N.G. and J.M. (Research Interns) are affiliated to Dr. Bagler’s lab at the Center for Computational Biology, and are thankful to IIIT-Delhi for the support. G.B. thanks Smita Sudheer, Resmi Vava, Vinay Randhawa and Shreyas Mangave for their feedback to improve the SpiceRx interface.

## Supplementary Information

### 1. Materials and Methods

The repertoire of culinary spices and herbs were collected from various sources including Foodb (http://foodb.ca/) Wikipedia <u>(https://en.wikipedia.org/wiki/List_of_culinary_herbs_and_spices), FPI (Food Plants International, http://foodplantsinternational.com/plants/)</u>, PFAF (Plants for a Future, http://www.pfaf.org) and FlavorDB (http://cosylab.iiitd.edu.in/flavordb) (Garg *et al.*, 2017). All 188 spices were normalized to NCBI Taxonomy IDs using their scientific names. These IDs were used to collect relevant abstracts of biomedical articles from MEDLINE in order to find empirical evidence for their disease associations.

MEDLINE article citations were downloaded from the FTP server of NCBI (ftp://ftp.ncbi.nlm.nih.gov/pubmed) and information regarding PMID, Date, Title, Abstract, Journal, and Authors was extracted using a modified version of PubMed parser (https://github.com/cosylabiiit/pubmed_parser). Named entity recognition was carried out in two stages. Spice/herb names in abstracts were tagged using a dictionary matching method. For recognizing and normalizing disease names in the text, we used TaggerOne (Leaman and Lu, 2016) which has a reported precision of 85% and recall of 80% on the Biocreative V Chemical Disease Relation test set (Li *et al.*, 2016). We used the pre-trained disease-only model available with TaggerOne (Leaman and Lu, 2016) on our data. Using TaggerOne, all disease names identified from abstracts were normalized to their corresponding MeSH IDs (Lipscomb, 2000).

We retained only those abstracts which had at least one spice and a disease mention. Sentence segmentation was performed on the retained abstracts to get candidate sentences for relation extraction. As a preprocessing step, all sentences which did not have at least one spice and a disease mention were removed. Further, to reduce the dimensionality of the word based feature space, we replaced all numbers with a standard identifier token and removed all special characters except punctuations. The sentences were then tokenized using GENIA (Kim *et al.*, 2003) and the part-of-speech (PoS) tag, as well as the chunk tag of each token were obtained.

To the best of our knowledge, no existing dataset exists for spice-disease or even food-disease associations. Thus, we manually annotated/labelled 6712 randomly selected sentences to tag positive, negative and neutral spice-disease associations. The training dataset consisted of 2669 sentences having positive associations and 301 sentences having negative association. Remaining sentences had neutral associations.

A Convolutional Neural Network (CNN) based relex model with with word (Mikolov *et al.*, 2013), position, PoS and chunk embedding features (Kumar Sahu *et al.*, 2016; Nguyen and Grishman, 2015) was trained on the labelled dataset. The word embedding was initialized using pre-trained weights from Chiu *et. al* (Chiu *et al.*, 2016) with the embedding for unknown words being initialized from a uniform (−*α*, *α*) distribution. The value of *α* was determined on the basis of the variance of embedding of known words. The position features consisted of distances of each token from the tagged entities. The embedding for PoS, position and chunk features was randomly initialized. Further, we zero-padded the sentences to equalize their lengths and concatenated different features of each token to form a single vector representation. The CNN consists of *n* layers. The first layer transforms the sentence into a *d* × *n* matrix, where *d* is the number of dimensions of the ‘token’ embedding and n is the length of the longest sentence in the corpus. The second layer consists of convolutions on the sentence embedding by *n*_*f*_ filters of filter size *f*_*z*_ with rectified linear unit (ReLU) activation function. The maximum activations from all filters are concatenated into a single vector of size *n*_*f*_ and fed to a Dropout layer (Srivastava *et al.*, 2014), which randomly sets an activation to zero with probability *p*. The next layer is a fully connected layer of *n*_*h*_ hidden units with ReLU activation function. Finally the output is fed to a softmax layer with 3 units. The categorical cross entropy loss was defined as the objective function and an _2_ regularization of 3 was applied on the fully connected layer. The network was trained using mini-batch gradient descent with shuffled batches of size 50 each and Adam (arXiv preprint arXiv: 1412.6980) optimizer. The model training was stopped if the validation loss did not decrease for 5 epochs. Oversampling of minority classes was used to address the problem of class imbalance, with the negative, positive classes being over-sampled by a factor of 12 and 1.35 respectively to the neutral class. The hyper-parameters of the neural network were determined by 5-fold cross validation.

After linking spices to diseases, we set out to explore the spice phytochemicals which might explain the molecular basis of the spice-disease associations. The phytochemical info for spices/herbs was obtained using KNApSAcK (Afendi *et al.*, 2012) and Comparative Toxicogenomics Database (CTD) (Davis *et al.*, 2017). All the compound identifiers were first standardized to PubChem and subsequently to MeSH in order to retrieve information regarding their bioactivity using PubChem BioAssay (Wang *et al.*, 2017) and therapeutic associations with diseases using CTD (Davis *et al.*, 2017).

### 2. Webserver Tech Stack

SpiceRx is implemented with the Python web development framework Django (https://www.djangoproject.com). and PostgreSQL (https://www.postgresql.org/). The frontend was built using HTML, CSS, JavaScript, AJAX, jQuery, JSME Molecular Editor (Bienfait and Ertl, 2013), Bootstrap, Jmol, DataTables and Google Charts. An Apache HTTP Server has been used to route requests to the Django application and to enable data compression for faster page load times. The site is best viewed in latest versions of Google Chrome, Firefox, Opera, Internet Explorer, and Microsoft Edge.

### 3. SpiceRx Statistics

Following are the statistics for data provided in SpiceRx.

#### 3.1 Statistics of research articles for spices/herbs

**Figure S1.**
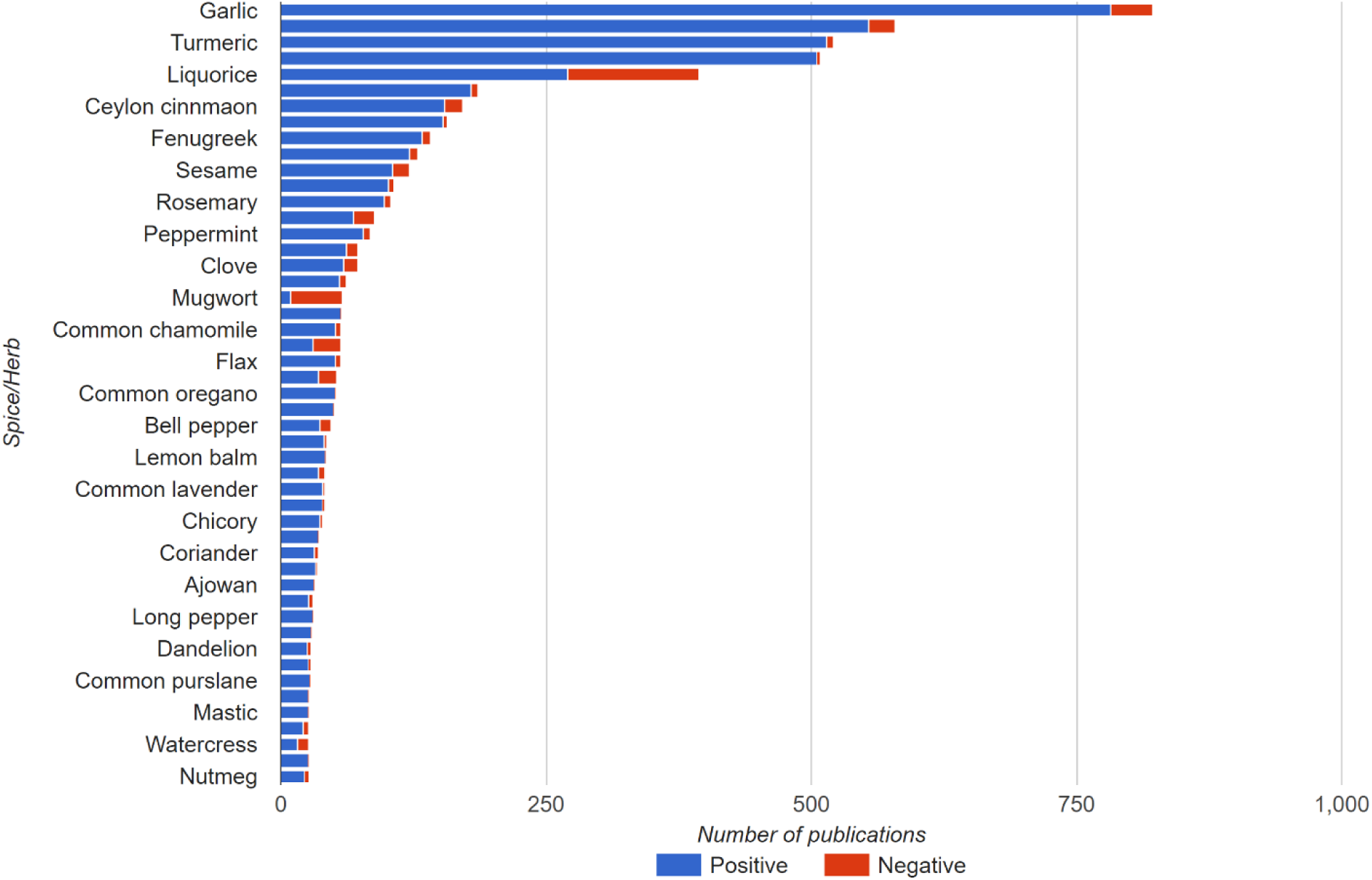
Statistics of research articles reporting positive and negative disease associations for spices/herbs with 10 or more publications. Statistics of research articles reporting positive and negative disease associations for Top 50 spices (with 10 or more publications). Among the spices/herbs with largest number of disease associations were Garlic (782+40), Ginkgo (554+25), Turmeric (515+6) and Ginger (505+3) with more than 500 research articles reported for each. Liquorice (123) and Mugwort (49) had the largest number of negative disease associations. For more details, please refer to the interactive graphics on the SpiceRx webpage: http://cosylab.iiitd.edu.in/spicerx/#stats

#### 3.2 Statistics of research articles for MeSH disease categories

**Figure S2.**
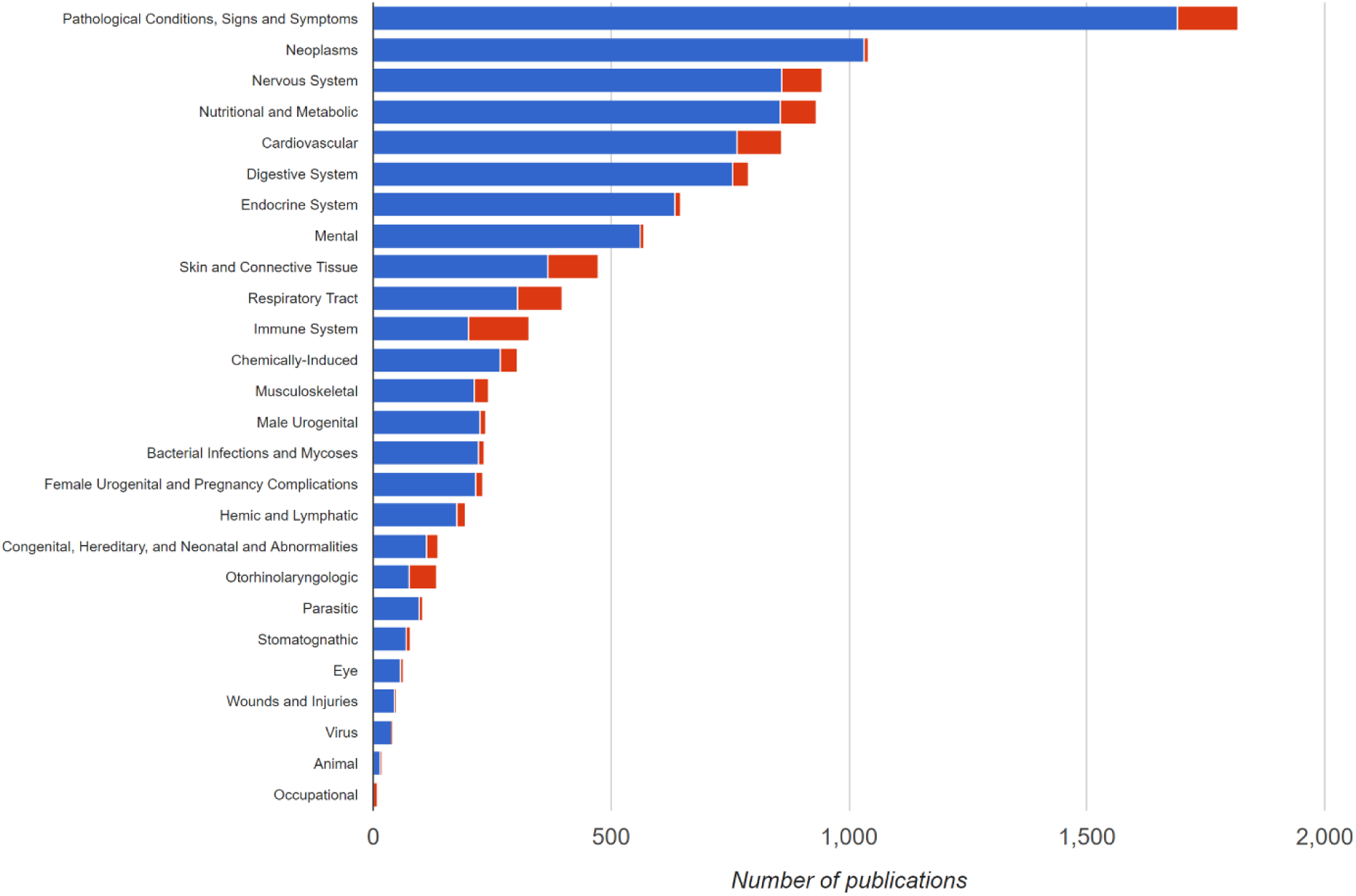
Statistics of research articles with reports of positive and negative associations of spices/herbs corresponding to MeSH disease categories. A total of 25 MeSH disease categories were reported with therapeutic and/or adverse effects for culinary spices and herbs. Among major disease categories that are influenced by spices/herbs were ‘Pathological conditions, Signs and Symptoms’ (1690+127), ‘Neoplasms’ (1033+9), ‘Nervous System Diseases’ (859+84), ‘Nutritional and Metabolic Diseases’ (857+74), ‘Cardiovascular Disease’ (765+95), ‘Digestive Systems Disease’ (755+35), ‘Endocrine Systems Diseases’ (635+10) and ‘Mental Disorders’ (561+7), with more than 500 research articles reported for each. Together, these MeSH categories cover a large number of diseases for which diet has been reported to have major role in pathological development and progression such as diabetes mellitus type 2, obesity, coronary artery disease, hypertension, and prostate cancer, among others. For more details, please refer to the interactive graphics on the SpiceRx webpage: http://cosylab.iiitd.edu.in/spicerx/#stats

#### 3.3 Statistics of research articles for therapeutic effects of phytochemicals

**Figure S3.**
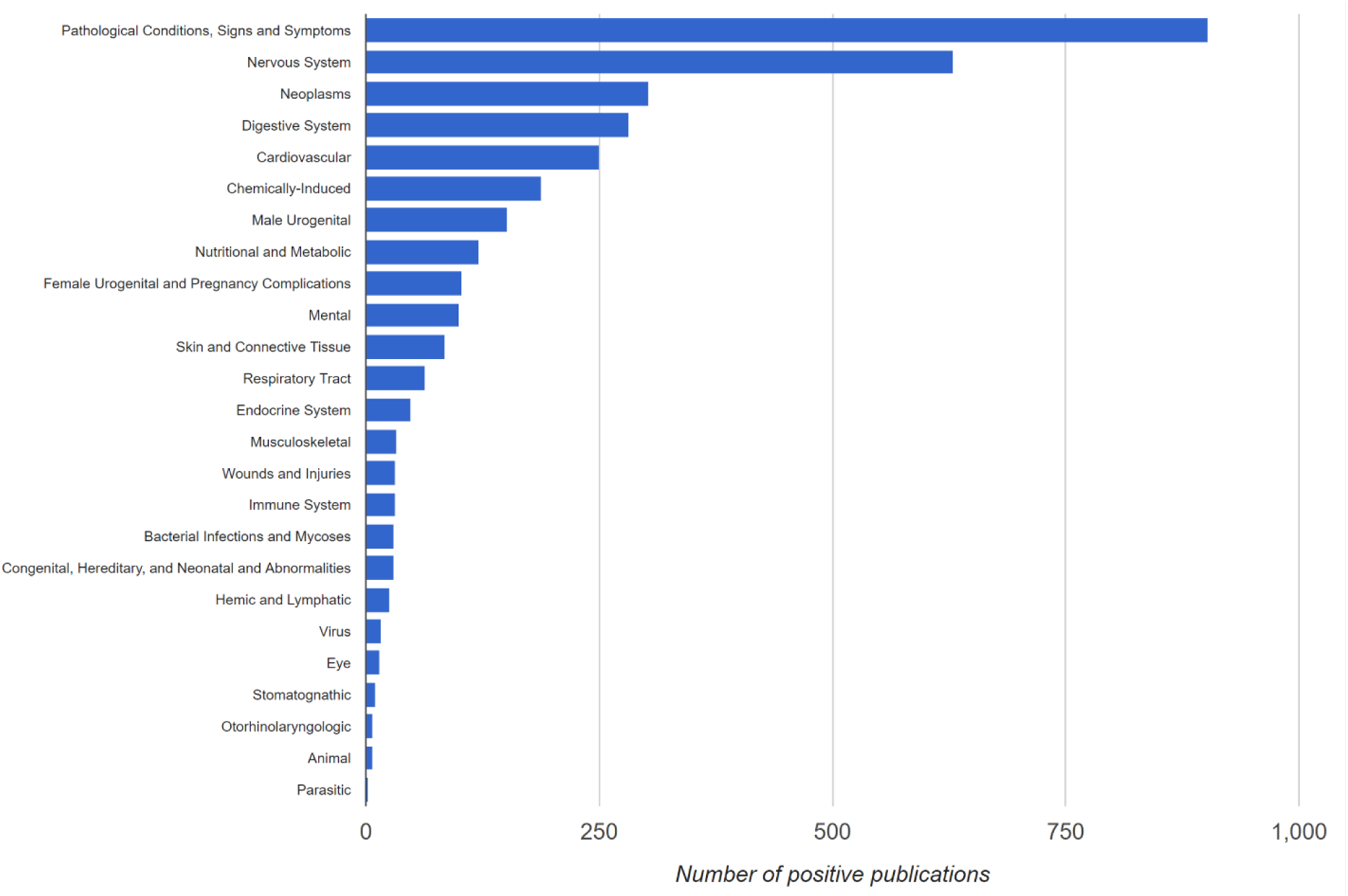
Statistics of research articles reporting positive associations for phytochemicals against MeSH disease categories. These data were obtained from Comparative Toxocogenomics Database. Amongst the top disease categories that phytochemicals were beneficial against were ‘Pathological conditions, signs and symptoms’ (903), ‘Nervous system disorders (629)‘, ‘Neoplasms’ (303) ‘Digestive system disorders’ (283) and ‘Cardiovascular disorders’ (250). To explore further, please refer to the interactive graphics on the SpiceRx webpage: http://cosylab.iiitd.edu.in/spicerx/#stats

#### 3.4 Molecular weight distribution of phytochemicals

**Figure S4.**
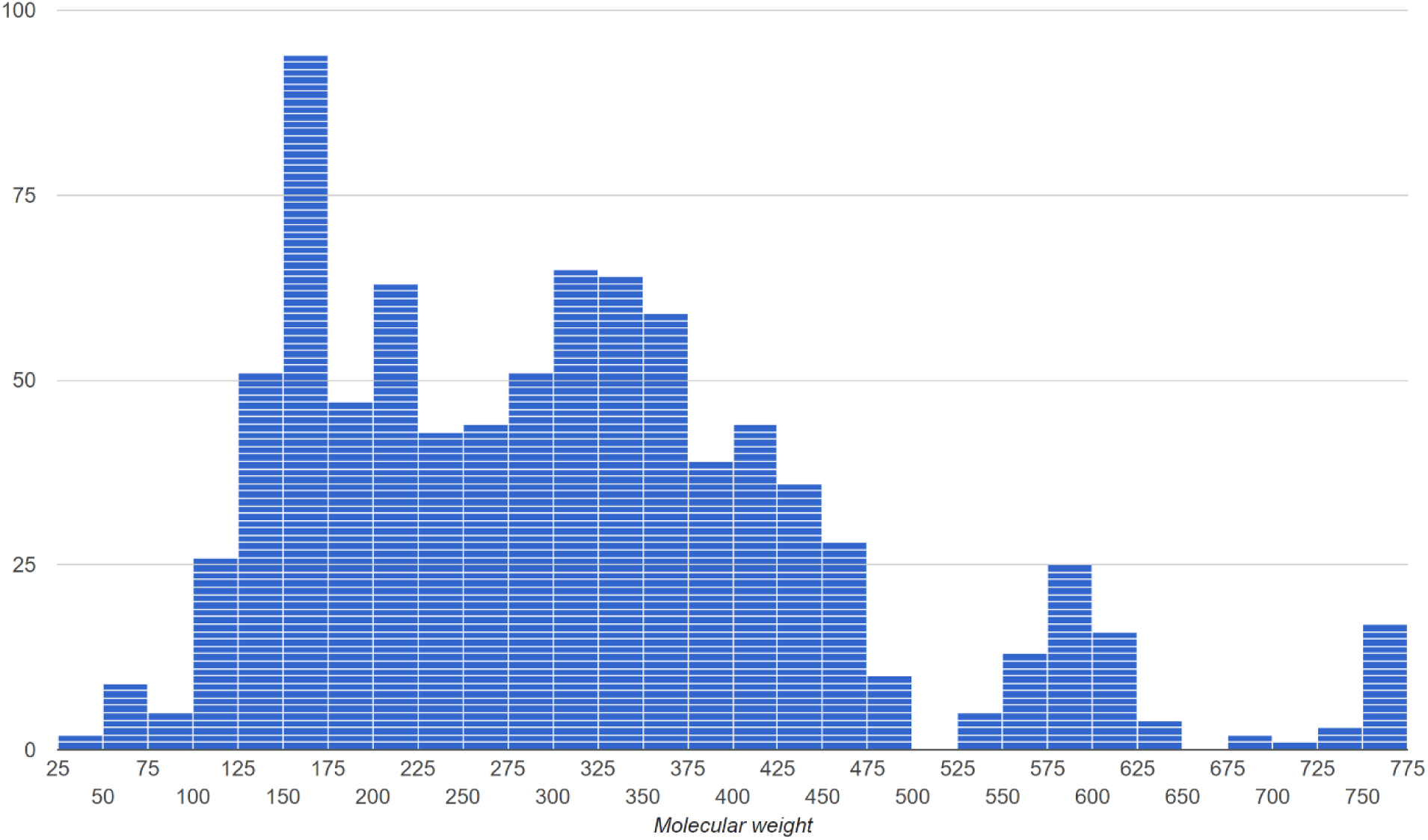
Distribution of molecular weights of phytochemicals from spices/herbs. Majority of the compounds had weight below 500 g/mol. To explore these data, please refer to the interactive graphics on the SpiceRx webpage: http://cosylab.iiitd.edu.in/spicerx/#stats

#### 3.5 Partition coefficient distribution of phytochemicals

**Figure S5.**
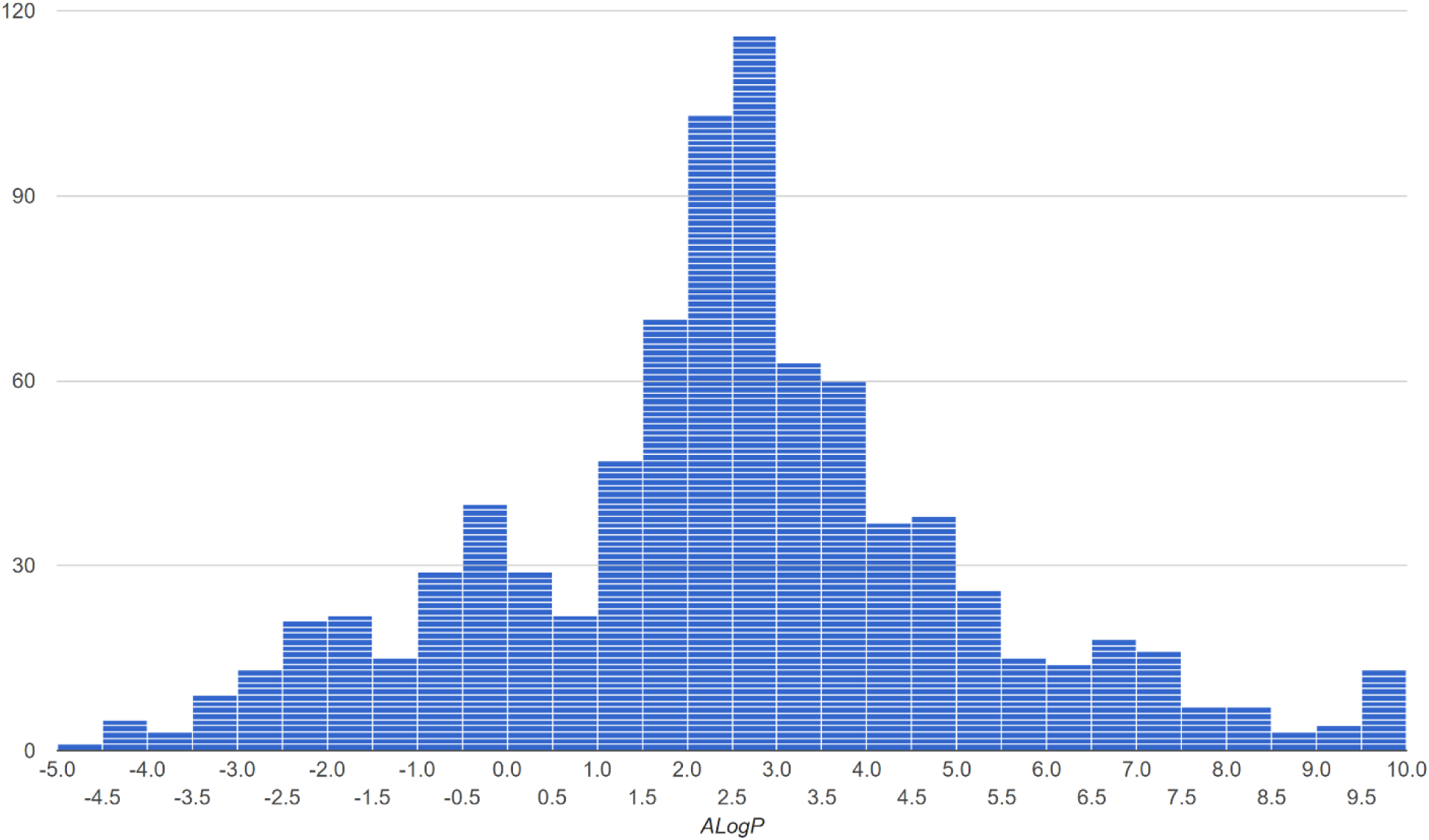
Distribution of partition coefficients (ALogP) of phytochemicals from spices/herbs. Majority of the molecules compounds had the partition coefficient below 5, thus fulfilling one of the criteria for drug-likeness. To explore these data, please refer to the interactive graphics on the SpiceRx webpage: http://cosylab.iiitd.edu.in/spicerx/#stats

### 4. Searching in SpiceRx

SpiceRx facilitates three types of searches— Search by Spices/Herbs, Diseases and Phytochemicals (Figure S6 and Figure S7).

- **Spice/Herb Search**: Users can search for the details of a Spice/herb in SpiceRx either using its common name, scientific name or the NCBI taxonomy ID (Figure S6(A)). For instance, one may search by either ‘saffron‘, ‘*Crocus Sativus*’ or ‘82528‘. As depicted in the illustrative figure, the user is assisted with autocomplete feature. The results page for a query displays the diseases associated with spice/herb (Figure S7(A)) and the spice phytochemicals.
- **Disease Search**: SpiceRx disease search options is aligned with the hierarchical organization of disease terms in MeSH (Figure S6(B)). Assisted with autocomplete, one may search by any disease name, disease category, disease subcategory or MeSH ID. To exemplify, the user can search by disease name ‘hypertension‘, its category, ‘cardiovascular diseases’ or subcategory, ‘vascular diseases‘. The disease search yields a page with details of spice/herb associations for the queried disease term (Figure S7(B)). Clicking on the ‘Details’ button presents with a list of spices associated with the disease as well as the phytochemicals involved in therapeutic association.
- **Phytochemicals Search**: SpiceRx may be searched for phytochemicals of spice/herb using the third option provided in the search panel (Figure S6(C)). Spice compounds may be searched using a range of molecular features such as its common name, MeSH ID, IUPAC name, PubChem ID, molecular weight, hydrogen bond donors/acceptors or molecular hydrophobicity (AlogP), apart from drawing their structure (JSME Molecular Editor (Bienfait and Ertl, 2013)). For instance, the users can search with common name ‘bisabolol‘, MeSH ID, ‘C004497‘, PubChem ID, ‘10586‘, IUPAC name, ‘6-methyl-2-(4-methylcyclohex-3-en-1-yl)hept-5-en-2-ol‘, Canonical SMILES, ‘CC1=CCC(CC1)C(C)(CCC=C(C)C)O‘, Molecular Formula, ‘C16H24O10’ or its Isomeric SMILES, ‘CC1=CCC(CC1)C(C)(CCC=C(C)C)O‘. The search generates a list of matching phytochemicals. Using the ‘Explore’ option against a phytochemical one can find a detailed characterization of the compound including details of spices in which the compound is reported, the diseases with which it is linked to, ADMET and physicochemical properties (Figure S7(C)). The 2D and 3D visualizations are provided using JSmol in addition to option for downloading in multiple formats. Importantly, one may search for similar molecules within SpiceRx of choose to find structurally similar molecules commercially available from ZINC (Irwin and Shoichet, 2005; Sterling and Irwin, 2015).

**Figure S6.**
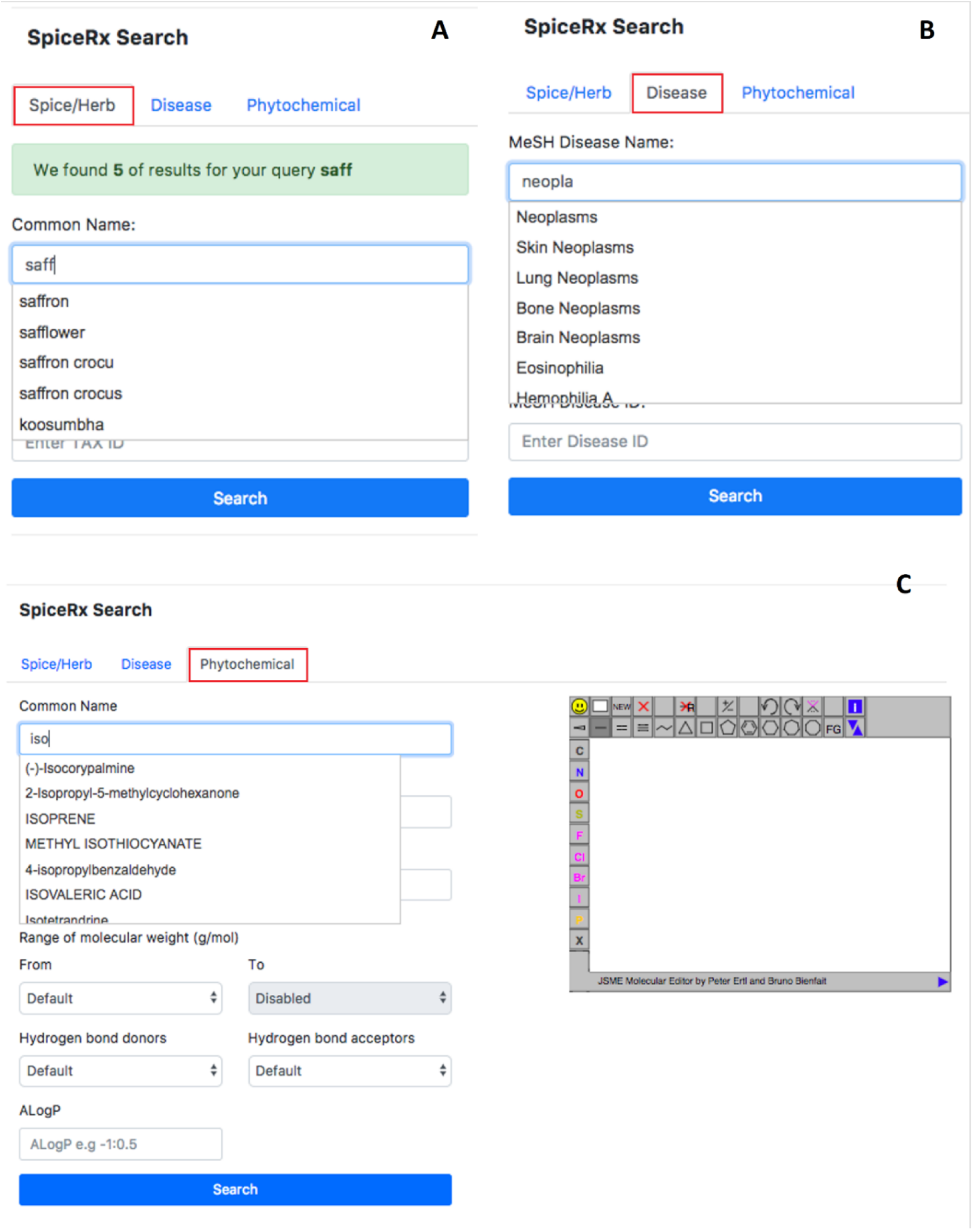
SpiceRx facilitates exploration of tripartite associations between spices/herbs, their phytochemicals and disease starting with any one of the elements: **A**. Spice, **B**. Disease, **C**. Phytochemical. Search panels are designed to enable experts and non-experts alike, and are assisted with multiple features such as autocomplete, spice synonyms, molecular drawing (JSME Molecular Editor) and such.

**Figure S7.**
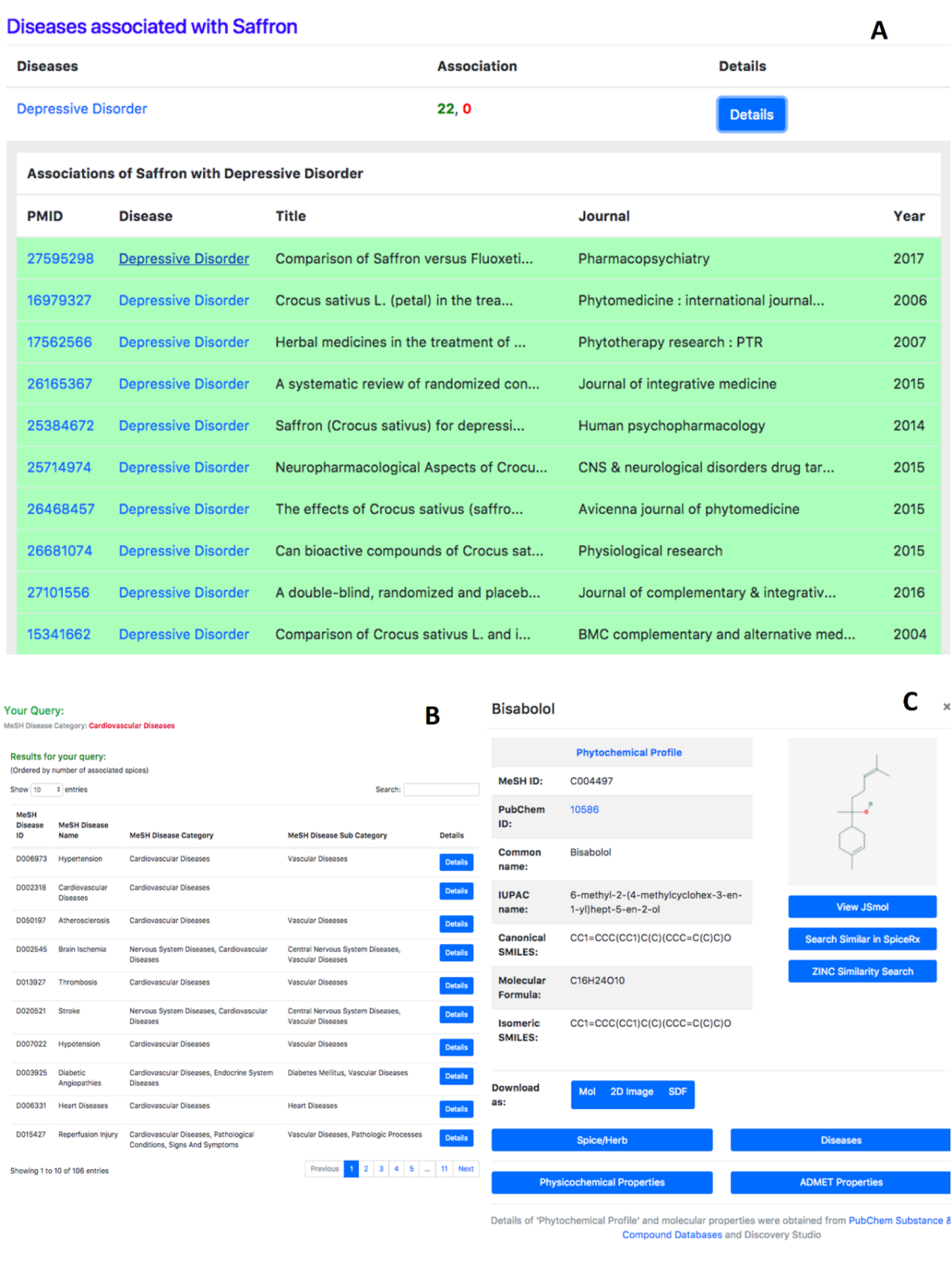
SpiceRx result panels for **A**. Spice Search, **B**. Disease Search, and **C**. Phytochemicals Explore panel. The user interface has been designed to enable discovery of interesting relationships through seamless exploration of triangular associations.

### 5. Case Studies

SpiceRx could be used for generating a variety of queries intended to find disease associations of spices and phytochemicals, to find spices linked to specific disease terms or for probing triangular associations between the three elements. Such queries could be aimed at finding ways for informed culinary use of spices and herbs against specific disease(s), discovering novel therapeutic spice-disease associations unreported hitherto, finding new ways of repurposing spices or finding potential spice phytochemicals against specific disease, among others. Here we present a few case studies.

#### (A) Searching for drug-like compounds in SpiceRx

One may search for spice phytochemicals which have drug-like properties using the phytochemical search tab in SpiceRx by using criteria for fulfilling Lipinski’s rule. Lipinski’s rule specifies conditions to evaluate drug-likeness of a compound; the suitability of a chemical compound to have pharmacological or biological activity making it a likely candidate for orally active drug. According to Lipinski’s rule, an orally active drug has no more than one violation of the following criteria: No more than 5 hydrogen bond donors; No more than 10 hydrogen bond acceptors; Molecular mass less than 500 Daltons; and an octanol-water partition coefficient (log P) not greater than 5.

**Figure S8.**
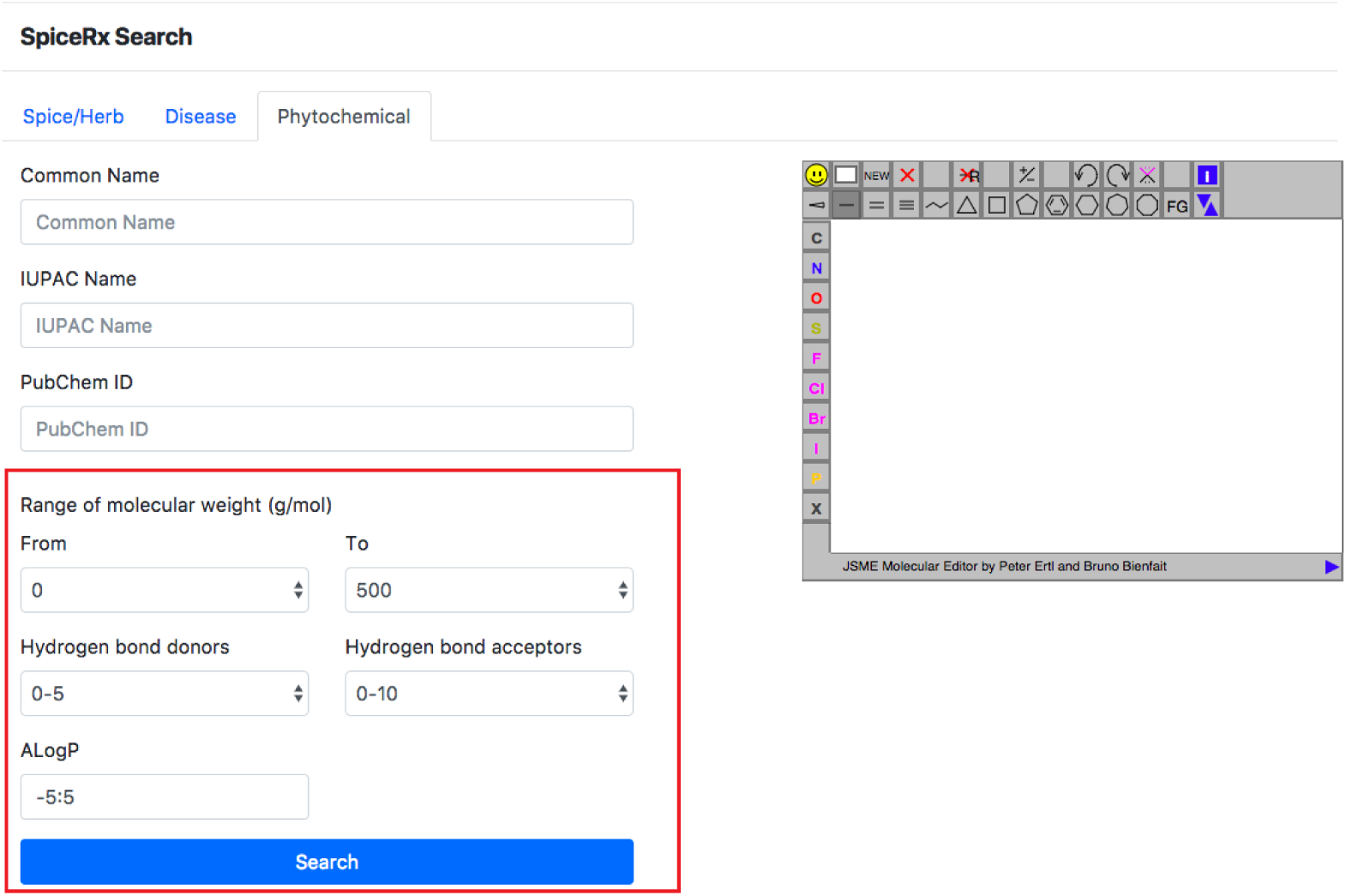
Drug-like phytochemicals in SpiceRx fulfilling Lipinki’s rule.

By specifying these conditions in the relevant search tabs, all compounds satisfying the above conditions can be obtained. This search in yields 639 (74 %) of all the phytochemicals in SpiceRx (866) suggesting the as potential therapeutic value of these compounds.

#### (B) Investigating molecular mechanisms behind therapeutic effects of spices and herbs

One of the key attributes of spices and herbs are that they are antimicrobial agents. Billing and Sherman (Billing and Sherman, 1998; Sherman and Billing, 1999) suggested that one of the reasons for the widespread use of spices in recipes is primarily due to their antimicrobial properties. One can search for spices and their phytochemicals which are active against microbial infections. Searching the broad category “Bacterial Infections and Mycosis” reveals that a wide range of culinary spices and herbs have positive associations against this category of diseases. Fennel and cinnamon have been reported with antimicrobial properties against salmonella infections. Common culinary spices and herbs such as garden thyme, oregano, clove, garlic, cinnamon, rosemary and turmeric have positive associations against “Pneumonia Staphylococcal”. Similarly, garlic, turmeric, oregano and clove are among spices which are reported to be beneficial for “*Escherichia coli* Infections”. Garlic clove and thyme are among spices which are reported to be beneficial for “Mycosis”. Interestingly, the molecular mechanism behind some of these antimicrobial effects of spices are yet to be uncovered. SpiceRx provides a fertile ground for further investigations into such cases where the molecular mechanisms for their actions are not yet evident.

Similarly, a large number of daily culinary spices and herbs have beneficial associations with Diabetes Mellitus, a common Nutritional and Metabolic Diseases. Among them, fenugreek, cinnamon, garlic, turmeric, ginger, and black cumin have been reported most frequently in literature. Beyond the three spice phytochemicals (’gallic acid‘, ‘berberine‘, ‘sophoraflavonoloside‘) which have been linked to diabetes mellitus, further investigations into the effects of phytochemical content in these spices can reveal their synergistic actions in disease regression and control.

#### (C) Hypothesis generation and knowledge discovery through SpiceRx

SpiceRx can be used as a platform for knowledge discovery and hypothesis generation. For specific diseases, SpiceRx provides phytochemicals which are therapeutically associated with them. By finding out the spices in which these phytochemicals occur, one can generate and test hypothesis confirming whether these spices have beneficial effects for these diseases. A few examples can further exemplify this point. ‘Glycyrrizin’ is a phytochemical therapeutically associated with ‘Hepatitis A‘. The phytochemical ‘Glycyrrizin’ is reported to be found in the herb liquorice. Similarly, ‘(-)-Epigallocatechin gallate’ is another phytochemical associated the disease ‘Herpes Simplex‘. This chemical is present in various spices such as peppermint, avocado leaf, bell pepper and german chamomile. These spices and herbs are not directly associated with Herpes Simplex which gives room for testing the hypothesis. Likewise, ‘Apigen’ is a phytochemical in many spices and herbs including thyme, coriander, hyssop and fennel among others, found to be effective for Adenoviridae Infections. No spices so far have been reported to be beneficial for Adenoviridae Infections. Further studies can be undertaken to confirm the efficacy of spices/herbs containing apigen against Adenoviridae Infections.

### 6. Data Download

All data are available for download: http://cosylab.iiitd.edu.in/spicerx/static/data/spicerx_dump.zip

### 7. Disclaimer

SpiceRx comes with the following disclaimer:

“All material on this website is a product of research and is provided for your information only and may not be construed as medical advice or instruction. No action or inaction should be taken based solely on the contents of this information; instead, readers should consult appropriate health professionals on any matter relating to their health and well-being.”

